# A Dynamical Model for the Low Efficiency of Induced Pluripotent Stem Cell Reprogramming

**DOI:** 10.1101/028266

**Authors:** Hussein Abdallah, Yili Qian, Domitilla Del Vecchio

**Affiliations:** Hussein Abdallah is with the Department of Electrical Engineering and Computer Science, Massachusetts Institute of Technology, 77 Mass. Ave, Cambridge MA; Yili Qian and Domitilla Del Vecchio are with the Department of Mechanical Engineering, Massachusetts Institute of Technology, 77 Mass. Ave, Cambridge MA

## Abstract

In the past decade, researchers have been able to obtain pluripotent stem cells directly from an organism’s differentiated cells through a process called cell reprogramming. This opens the way to potentially groundbreaking applications in regenerative and personalized medicine, in which ill patients could use self-derived induced pluripotent stem (iPS) cells where needed. While the process of reprogramming has been shown to be possible, its efficiency remains so low after almost ten years since its conception as to render its applicability limited to laboratory research. In this paper, we study a mathematical model of the core transcriptional circuitry among a set of key transcription factors, which is thought to determine the switch among pluripotent and differentiated cell types. By employing standard tools from dynamical systems theory, we analyze the effects on the system’s dynamics of overexpressing the core factors, which is what is performed during the reprogramming process. We demonstrate that the structure of the system is such that it can render the switch from an initial stable steady state (differentiated cell type) to the desired stable steady state (pluripotent cell type) highly unlikely. This finding provides insights into a possible reason for the low efficiency of current reprogramming approaches.

## I. INTRODUCTION

Stem cells are unspecialized precursors with the potential to differentiate into more specialized, differentiated cell types. In particular, embryonic stem cells (ESCs) are at the top of the differentiation hierarchy and can give rise to any cell in an organism [1]. This makes them popular candidates for use in regenerative and therapeutic medicine, where they can replace a patient’s injured or diseased cell type of choice [2]–[4]. However, ESC therapies face a couple of major limitations. ESCs derived from early fetal cell masses confront various ethical controversies [5], and transplanting another organism’s stem cells into a patient runs the risk of failed engraftment due to the patient’s immunogenic barrier [6].

The biological and ethical factors limiting the efficacy of ESCs make iPS cells a much more preferred alternative, since they face neither of these issues by their very nature. First discovered in a seminal experiment by Yamanaka et. al in 2006, iPS cells were produced by overexpressing only four pluripotency-related transcription factors in differentiated cells [7]. Since then, this strategy has been used to create a variety of iPS cells from various differentiated cell types [8]. However, there is still a large bottleneck in iPS cell research due to the low efficiency of reprogramming in these experiments, generally around 1% or much less [9]–[11].

In this paper, we investigate structural aspects of these overexpression-based reprogramming strategies to explain potential reasons for this failure. Specifically, we study the dynamics of three transcription factors–Oct4, Sox2, and Nanog (OSN)–which have emerged as the master regulatory factors involved in pluripotency [12]–[18]. Specific levels of OSN characterize three cell types, each represented by a stable steady state (SSS) of the dynamic model for OSN. These include embryonic stem cells (which are pluripotent) and their more differentiated counterparts, the trophectoderm (TR) and primitive endoderm (PrE) cells (which are multi-potent).

Our dynamical systems approach is related to the well-established Waddington epigenetic landscape metaphor [19]. In the Waddington framework, phenotypical cell types are represented as basins of attraction in a landscape of equilibria. In this model, biological cell differentiation is understood as a transition between attractors’ basins due to stimuli applied to the underlying genetic network [20]. In our specific case, the ESC, TR, and PrE states are basins of attraction, and the reprogramming process aims at transforming a state in the TR (or PrE) basin of attraction to a state in the ESC basin of attraction.

This paper is organized as follows. In Section II, we begin with a 3D ordinary differential equation (ODE) model for the dynamics of OSN, which we reduce to a 2D ODE model between Oct4 and Nanog. We then use nullcline analysis to characterize the ESC, TR, and PrE cell types as stable steady states (SSS) in the 2D model. In Section III, we investigate the dynamics of overexpression-based reprogramming. We discover that current overexpression strategies rarely succeed in reprogramming differentiated cells (in this case, TR or PrE cells) into pluripotent cells (ESCs, or iPS cells), due to the nullclines’ structure.

## II. MODEL

### A. The Oct4-Sox2-Nanog Fully Connected Triad

Fig. 1 summarizes the early embryological steps a zygote undergoes after fertilization to divide and differentiate into cells with decreasing plasticity. These are the differentiation steps which give rise to the three cell types of interest: ESCs, TRs, and PrEs. The working hypothesis of the identity of the molecular machinery dictating which cells go down each path in either differentiation step is the Oct4-Sox2Nanog network [12]–[18]. We approximate the topology of this network as a fully connected triad (FCT) among Oct4, Sox2, and Nanog, as shown in Fig. 2. FCTs are a class of 3-node network motifs in which all nodes directly interact with one another. In particular, when these interactions are all activations, these special FCTs have been shown to exhibit the desired multi-stability properties we seek [21]. The FCT we model here contains an additional repression mechanism, which has been more recently suggested [22].

**Fig. 1:**
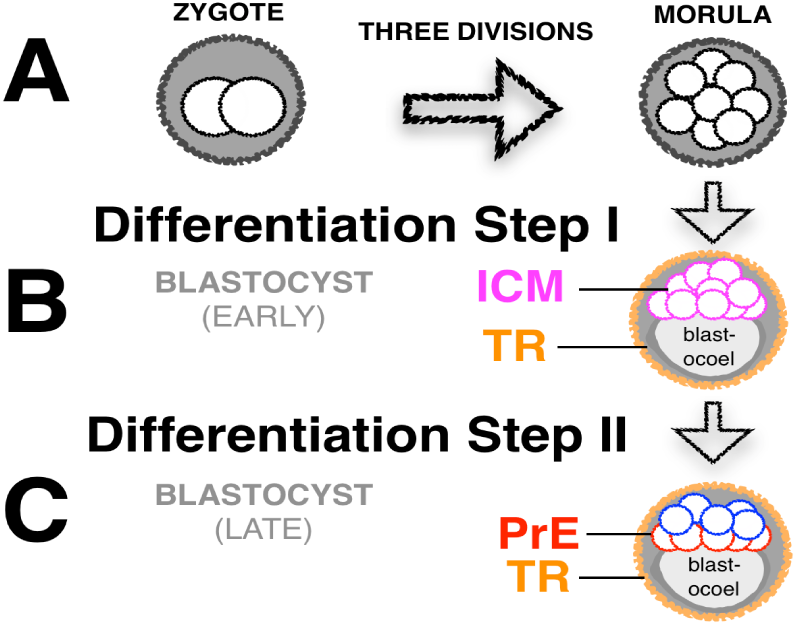
(A) In the first steps after fertilization, three rounds of cell division form an 8-cell mass known as the morula. (B) The first differentiation event occurs when an epithelial layer known as the trophectoderm (TR) forms around the perimeter of the mass, leaving the rest of the cells in an inner cell mass (ICM), (from which ESCs are derived [23], [24]), pushed against the blastocoel cavity. (C) The second differentiation event occurs when ICM cells adjacent to the blastocoel form an epithelium known as the primitive endoderm (PrE), leaving the rest of the cells to the epiblast lineage. [25] was used in creating this figure.

**Fig. 2:**
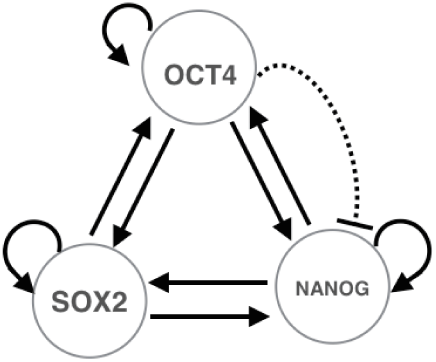
The network of Oct4, Sox2, and Nanog modeled as a fully connected triad (FCT).

### B. The Nonlinear 3D OSN Model

Referring to the topology of Fig. 2, the following ordinary differential equation model based on cooperative Hill functions [26] is employed to describe the dynamics of the FCT. Let *N, O* and *S* represent the concentrations of Nanog, Oct4, and Sox2, respectively. The model is given by

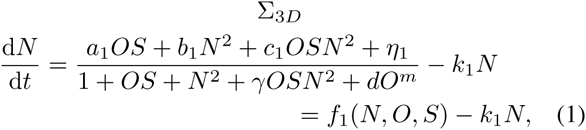

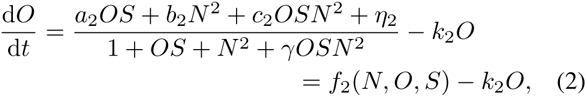

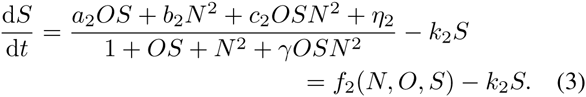

In the ODEs above, the linear terms are due to dilution and degradation, and all parameters are positive. The nonlinear Hill function terms, denoted by *f_i_* (*i*=1,2), model the species’ interaction with one another. Specifically, the Hill functions incorporate activation by the heterodimer *OS*, the homodimer *N*^2^, and the molecule *OSN*^2^. Additional features and assumptions of model Σ_3_*_D_* are described below.

1. *Nanog Dimerization:* In contrast to the models of [27], [28], which consider Nanog as a monomer, it is treated as a homodimer (*N*_2_) in Σ_3_*_D_*. This is based on strong evidence suggesting Nanog only binds to other pluripotency factors when dimerized [29], [30].
2. *Independent Nanog Promoter Activity:* The models of [27], [28] only consider activation by Nanog when this species is bound to the Oct4-Sox2 heterodimer. While it has been shown that binding Oct4-Sox2 significantly strengthens activation by Nanog [31], the species can still act as an activator – albeit a much weaker one – when not bound to the heterodimer. In [31], activation by Nanog when one or both of Oct4 and Sox2 were mutated was reduced to between 6-17% of wild type activity. To capture this, independent *N*_2_ activator terms are included in the Hill functions of Σ_3_*_D_*. We note the rough quantitative constraint on the relative strength of this term to the *OSN*_2_ term when determining appropriate values for the *b_i_* and *c_i_* parameters in sections below.
3. *Repression of Nanog by Oct4:* In [22], it was reported that low levels of Oct4 were correlated with high Nanog and higher levels of Oct4 were correlated with low Nanog. Although no regulatory link was demonstrated, we model this empirical observation with a higher order repressive term by Oct4 on Nanog, (*dO_m_*,*m* > 2), as seen in (1). Based on evidence suggesting Oct4 forms a heterodimer with Sox2 [32], we require that the order of repression is an even number.
4. *Equal Dynamics for Oct4 and Sox2:* Given that Oct4 and Sox2 are known to work together [33] and have been considered as the same species in previous models [22], here we assume that their dynamics are the same. This simplifies the analysis without affecting the main conclusions.
5. *Denominator Constants Normalized to 1:* The standard form of a Hill function for an activator *x_a_* is *a*_1_(*x_a_/k_a_*)*_n_*/(1 + *b*_1_(*x_a_/k_a_*)*_n_*), and for a repressor *x_r_* it is 1*/*(1 + *b*_1_(*x_r_/k_r_*)*_n_*). As seen in (1)-(3), the parameters in the denominators of the Hill functions are set to 1 for every activator and repressor. This is performed via normalization as follows. Given a Hill function of the form

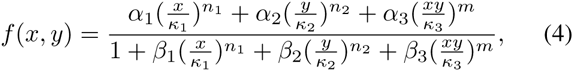

the substitutions 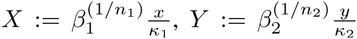 allow (4) to be rewritten as

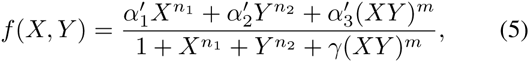

where 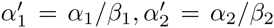, 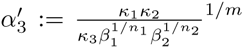, 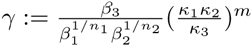.

### C. Model Reduction into a 2D System

To study the location and stability of the steady states of Σ_3_*_D_*, we study a reduced order system given by

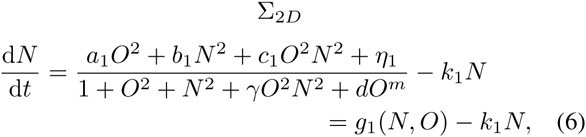

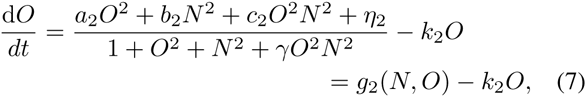

which is obtained by substituting *S* = *O* in (1) and (2). We justify this reduction with the claims below, which show that the location and stability of steady states of Σ_3_*_D_* can be studied by studying those of Σ_2_*_D_*.

#### Lemma

If (*N*^∗^,*O*^∗^, *S*^∗^) is a steady state of Σ_3_*_D_*, then *S*^∗^ = *O*^∗^.

##### Proof

Stable states (*N*^∗^, *O*^∗^, *S*^∗^) of Σ_3_*_D_* satisfy:

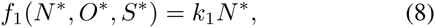

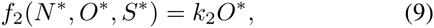

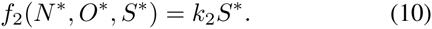

Substitution of (10) into (9) shows *S*^∗^ = *O*^∗^■

Any steady states of Σ_3_*_D_* are assumed to be of the form (*N*^∗^,*O*^∗^, *O*^∗^) in what follows. The claim below shows that every steady state in Σ_3_*_D_* maps to a unique steady state in Σ_2_*_D_*. Conversely, any steady state of Σ_2_*_D_* maps to a unique steady state in Σ_3_*_D_*.

#### Claim 1

(*N*^∗^, *O*^∗^, *O*^∗^) is a steady state of Σ_3_*_D_* if and only if (*N*^∗^, *O*^∗^) is a steady state of Σ_2_*_D_*.

##### Proof

(⇒) If (*N*^∗^, *O*^∗^, *O*^∗^) is a steady state of Σ_3_*_D_*, then *f*_1_(*N*^∗^, *O*^∗^, *S*^∗^)= *k*_1_*N*^∗^, *f*_2_(*N*^∗^, *O*^∗^, *S*^∗^)= *k*_2_*N*^∗^. Since

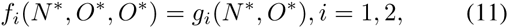

then we also have *g*_1_(*N*^∗^, *O*^∗^)= *k*_1_*N*^∗^ and *g*_2_(*N*^∗^, *O*^∗^) = *k*_2_*O*^∗^. This implies the dynamics of Σ_2_*_D_* satisfy 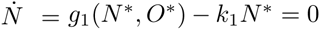 and 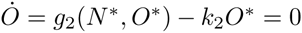. Therefore, (*N*^∗^, *O*^∗^) is a steady state of Σ_2_*_D_*.

(⇐) If (*N*^∗^, *O*^∗^) is a steady state of Σ_2_*_D_*, then *g*_1_(*N*^∗^, *O*^∗^)= *k*_1_*N*^∗^, *g*_2_(*N*^∗^, *O*^∗^)= *k*_2_*O*^∗^. Due to (11), we also have *f*_1_(*N*^∗^, *O*^∗^, *O*^∗^)= *g*_1_(*N*^∗^, *O*^∗^)= *k*_1_*N*^∗^ *f*_2_(*N*^∗^, *O*^∗^, *O*^∗^) = *g*_2_(*N*^∗^, *O*^∗^)= *k*_2_*O*^∗^ = *k*_2_*S*^∗^, which implies (*N*^∗^, *O*^∗^, *O*^∗^) is a steady state of Σ_3_*_D_.*■

In addition, the claim below shows that the stability properties of a steady state in Σ_2_*_D_* reflect the same properties for the corresponding steady state in Σ_3_*_D_*.

#### Claim 2

(*N*^∗^, *O*^∗^) is a stable (unstable) steady state of Σ_2_*_D_* if and only if (*N*^∗^, *O*^∗^, *O*^∗^) is a stable (unstable) steady state of Σ_3_*_D_.*

##### Proof

We first show that 2D stability implies 3D stability. The Jacobians of systems Σ_2_*_D_* and Σ_3_*_D_* are given by

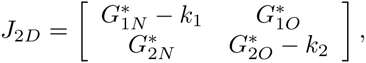

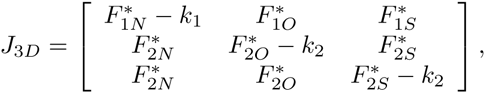

respectively, where 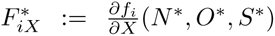, 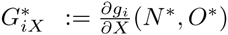. Stability of a stable state in Σ_2_*_D_* implies:

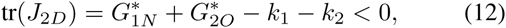

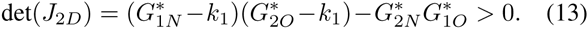

The characteristic equation of Σ_3_*_D_* can be written as:

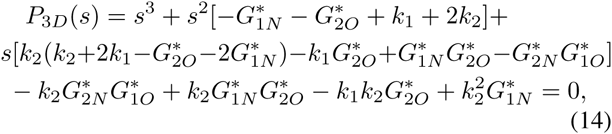

where the relation

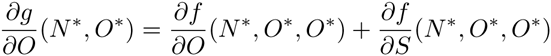

is used, which comes from using the chain rule on *g*(*N, O*) = *f*(*N, O, z*(*O*)), *z*(*O*) = *O*. Equation (14) can be rewritten to obtain

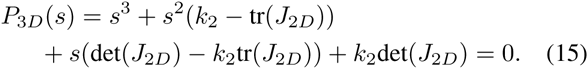

The Routh-Hurwitz stability criterion states that a third-order system with characteristic polynomial *P*(*s*)= *a*_3_*s*^3^+*a*_2_*s*^2^+ *a*_1_*s* + *a*_0_ is stable if and only if *a_n_* > 0 for all *n* and *a*_2_*a*_1_ *> a*_3_*a*_0_. As a consequence, the stability conditions for Σ_3_*_D_* become:

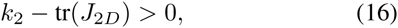

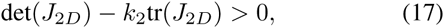

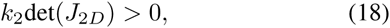

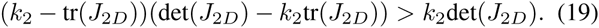

Relations (16)-(18) are trivially implied by the stability conditions of Σ_2_*_D_*, while (19) can be reduced to:

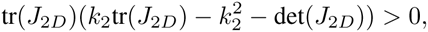

which, in turn, is also implied by the stability conditions (12)-(13) of Σ_2_*_D_* since *k_i_* > 0. This shows that stability of (*N*^∗^, *O*^∗^) in Σ_2_*_D_* implies stability of (*N*^∗^, *O*^∗^, *O*^∗^) in Σ_3_*_D_*. We now show that instability of (*N*^∗^, *O*^∗^) in Σ_2_*_D_* implies instability of (*N*^∗^, *O*^∗^, *O*^∗^) in Σ_3_*_D_*. The eigenvalues of Σ_2_*_D_* are

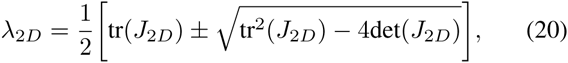

and there are two cases in which any of these are positive: i) det(*J*_2_*_D_*) < 0 and ii) det(*J*_2_*_D_*) > 0, tr(*J*_2_*_D_*) > 0. If (*N*^∗^, *O*^∗^) in Σ_2_*_D_* is unstable and we are in case i), (18) is easily seen to be not satisfied, making Σ_3_*_D_* unstable. If we are in case ii), it can be shown that conditions (16)–(19) cannot simultaneously be satisfied. Without loss of generality, we can assume (16)-(18) are satisfied, and test if condition (19) is satisfied. It is satisfied if

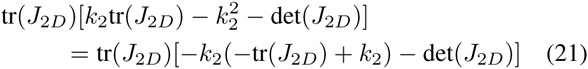

is positive. The right hand side of (21) is negative since tr(*J*_2_*_D_*) > 0, det(*J*_2_*_D_*) *>* 0 and −*k*_2_(−tr(*J*_2_*_D_*)+ *k*_2_) < 0 from assuming (16)-(18) is true. This shows that instability of (*N*^∗^, *O*^∗^) in Σ_2_*_D_*.■

By virtue of these claims, we can focus on studying the number and stability of the steady states of Σ_2_*_D_*.

### D. Characterizing ESC, Tr, PrE States by Oct4 and Nanog Levels

Using experimental studies of the differentiation steps that give rise to the ESC, TR, and PrE states, Table 1 summarizes the results of a literature search concluding with the characterizations of the three cell types shown in Fig. 3. While the experiments show that Nanog levels in ESC are much higher than those in TR and PrE, we found no experimental evidence comparing Nanog levels in TR and PrE. We assume that Nanog in PrE is at an intermediate level between Nanog in TR and ESC. There is no loss of generality because the strength of the parameter d in (6) can be chosen to further decrease Nanog levels for high Oct4. With this characterization and the mathematical framework established above, we now form a correspondence between the SSS of Σ_2_*_D_* and the physiological cell types corresponding to certain levels of Oct4 and Nanog.

**TABLE I:** Results of literature search characterizing Oct4 & Nanog levels in TR, ESC, PrE cell states

|  |  |  |
| --- | --- | --- |
| **Oct4** | **TR <ESC** | **ESC <PrE** |
| **Levels** | [34]–[37] | [36]–[38] |
| **Nanog** | **TR, PrE <ESC** |  |
| **Levels** | [14], [18], [39] |  |

**Fig. 3:**
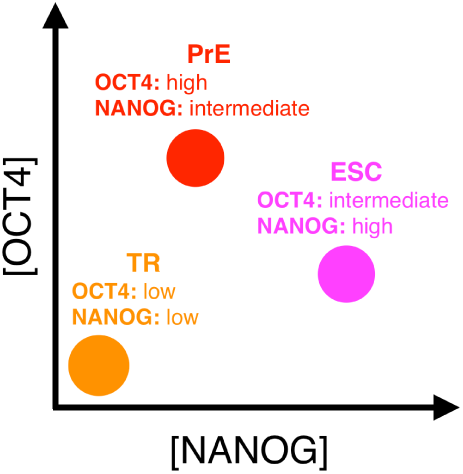
State characterizations of ESC, TR, and PrE based on relative Oct4 and Nanog concentrations determined from Table 1.

### E. 2D Nullcline Analysis

We perform nullcline analysis to determine the SSS corresponding to the qualitative levels of Oct4 and Nanog in each cell type seen in Fig. 3. The nullclines describing the mathematical steady states of Σ_2_*_D_* at equilibrium 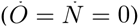 are

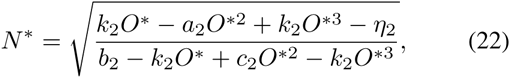

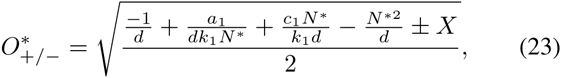

where we have defined

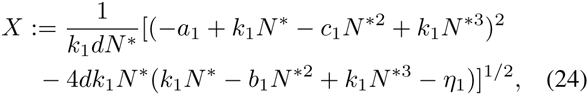

and have set the order of Oct4 repression on Nanog to *m =*4 for the sake of analysis.

The main parametric constraint used when performing this analysis was that the strength, *b_i_*, of activator *N*_2_ should be less than the strength, *c_i_*, of *OSN*_2_–on the order of about 0.10. This is in accordance with the experimental report [31] showing Nanog promoter activity when Oct4 and Sox2 were mutated was measured to be much less than wild type activity (6-17%). In Fig. 4A, nullclines are shown for a representative parameter set which satisfies these constraints (*b*_1_/*c*_1_ ≈ 0.27, *b*_2_/*c*_2_ ≈ 0.09). There are five steady states, whose stability is studied through geometric reasoning as follows.

**Fig. 4:**
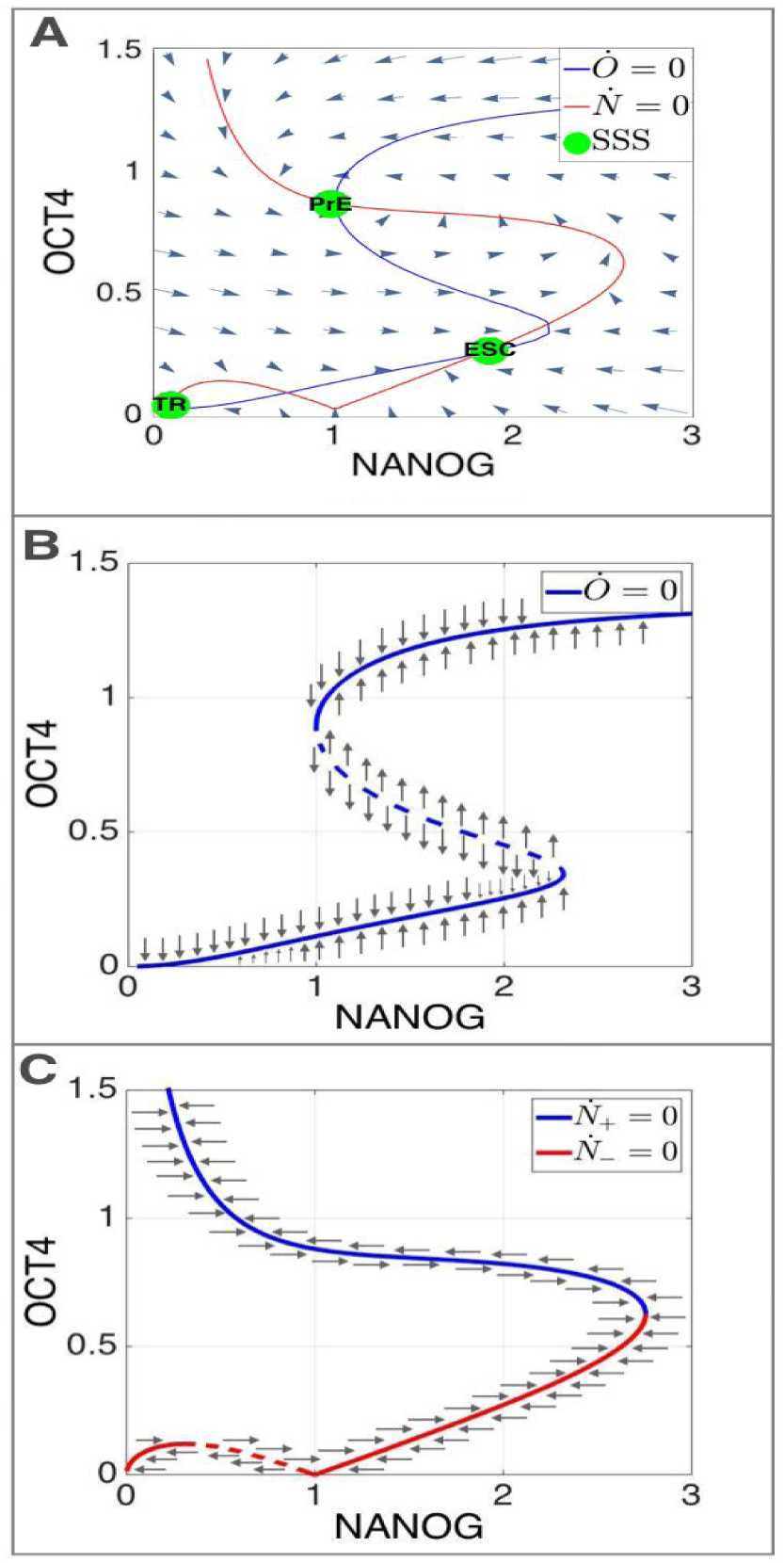
Nullcline stability analysis: (A) nullclines for the parameter set giving rise to 3 SSS: *m* =4, *a*_1_ = 10, *b*_1_ = 2, *c*_1_ = 7.5, *k*_1_ = 1, *η*_1_ = 0.0001, *a*_2_ =1.8, *b*_2_ =0.18, *c*_2_ =2, *k*_2_ = 1, *η*_2_ = 0.0001, *γ* = 1, d = 20. *d* is chosen to reduce Nanog levels in PrE (high Oct4) so they remain under levels in ESC. The vector field is plotted using Mathematica V10.0.2.0. (B) Sign of the vector field in the O direction in the proximity of the nullcline 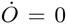. (C) Sign of the vector field in the *N* direction in the proximity of the nullcline 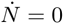.

We define *H*_1_(*N, O*):= *g*_1_(*N, O*) − *k*_1_*N, H*_2_(*N, O*):= *g*_2_(*N, O*)−*k*_2_*O*. We first study the stability of a point (*N, O*) on the nullcline *H*_2_(*N, O*) = 0 (Fig. 4B) in the *O* direction. For a fixed *N*, the corresponding point on the nullcine is stable in the *O* direction if *∂H*_2_*/∂O* < 0. Using the implicit function theorem, we have

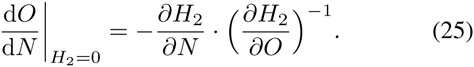

Graphically, the left hand side in (25) represents the local slope of the nullcline in Fig. 4B. In our system, *∂H*_2_*/∂N* > 0 for all positive (*N, O*). Therefore, *∂H*_2_*/∂O* < 0 if and only if d*O/*d*N* > 0. As a result, a point (*N, O*) on the nullcline *H*_2_(*N, O*) = 0 is stable in the O direction if and only if d*O/*d*N* is positive at (*N, O*).

Similarly, when studying the stability of a point (*N, O*) on *H*_1_(*N, O*) = 0 in the *N* direction, we consider the nullcline d*N/*d*t* = *H*_1_(*N, O*) = 0 in Fig. 4C. For a fixed *O*, the steady state is stable in the *N* direction if *∂H*_1_*/∂N* < 0. Applying the implicit function theorem, we obtain the condition that the steady state is stable if and only if

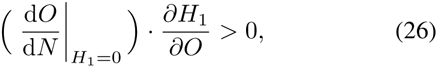

where the first term in (26) is the local slope of the null-cline in Fig. 4C. In contrast to the previous case, however, *∂H*_1_*/∂O* is not sign definite when evaluated on the nullcline *H*_1_(*N, O*) = 0. To understand which segment of the null-cline in Fig. 4C has positive (negative) *∂H*_1_*/∂O*, we note that for a fixed *N* = *N*^∗^, *O* is solved by a quadratic function of *O*_2_,

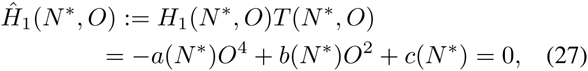

where *a* = −*k*_1_*dN*^∗^, *b* = *a*_1_ − *k*_1_*N*^∗^ + *c*_1_*N*^∗2^ − *k*_1_*N*^∗3^, *c* = −*k*_1_*N*^∗^ − *k*_1_*N*^∗3^ + *b*_1_*N*^∗2^ + *η*_1_ and *T* (*N*^∗^,O):= 1+ *O*^2^ + *N*^∗2^ + *O*^2^*N*^∗2^ > 0 is the denominator term in *g*_1_(*N*^∗^, *O*^∗^). When (27) has two solutions 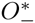 (red branch in Fig. 4C) and 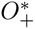 (blue branch in Fig. 4C), we have 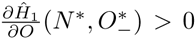 and 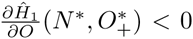. Moreover, since 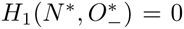 at 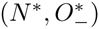,

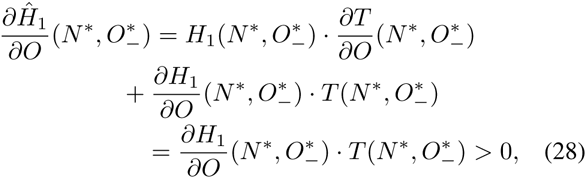

which implies that *∂H*_1_*/∂O* > 0 at 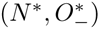. Similarly, we can show that *∂H*_1_*/∂O* < 0 at 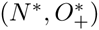. Therefore, the red branch (corresponding to 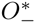) is stable (unstable) in the *N* direction when the local slope of the nullcline in Fig. 4C is positive (negative). Conversely, the blue branch (corresponding to 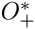) is stable in the *N* direction when the nullcline has negative local slope, and unstable otherwise. Given the above conclusions, since the system is in 2D, a steady state (*N*^∗^, *O*^∗^) is stable if and only if it is stable in both the *O* and *N* directions. According to this stability analysis, the nullclines in Fig 4. lead to 3 SSS and 2 unstable steady states.

## III. Reprogramming Using Overexpression

In a 2006 report by Yamanaka and colleagues, a small cocktail of only 4 transcription factors (among which were Oct4 and Sox2) was identified as the sole requirement for reprogramming differentiated cells into stem cells [7]. In this experiment, adult fibroblast cells were induced into pluripotent stem cells using the transcription factor cocktail. Since then, a variety [8] of differentiated cells and experimental conditions have been used to produce iPSCs, yet the efficiency of reprogramming remains around 1% or much less [9]–[11].

Here we use nullcline analysis to provide a possible explanation as to why current techniques show this systematic failure after nearly 10 years of practice. In overexpression experiments, transcription factors are produced in excess in a cell using viral and non-viral vectors [40]. Specifically, the genes encoding the transcription factors are inserted into these vectors under the control of constitutive or inducible promoters. This effectively increases the production rate of the factors. In our model, we therefore treat this overexpression as an additional constant term added to the production rate of the factors. This leads to the modified system:

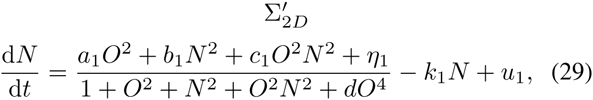

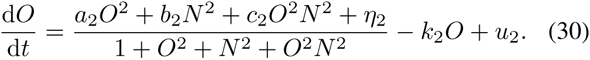

in which *u*_1_ and *u*_2_ are positive and can be effectively viewed as inputs to the system. The nullclines change to

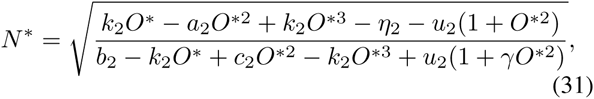

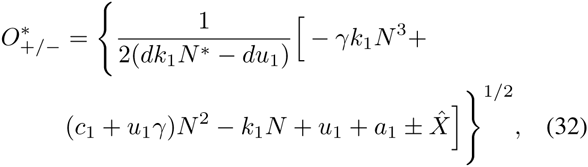

where we have defined

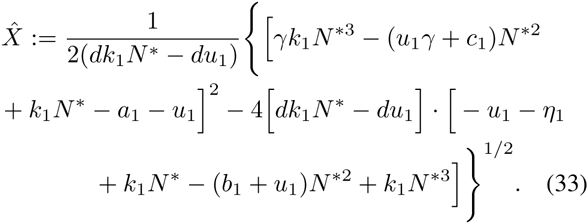

We use these nullclines to demonstrate the potential problem in seeking to force transitions from a differentiated state to the ESC state by constant inputs *u*_1_ and *u*_2_. In particular, we start with modeling reprogramming experiments with only the overexpression of Oct4 (employing *u*_2_ *>* 0*, u*_1_ = 0), which has been used to produce iPS cells on its own from various differentiated cells [41]–[46]. We specifically model an experiment reprogramming the TR state to ESC using only Oct4, as in [46]. For the sake of completeness, we then consider reprogramming experiments with only Nanog overexpression (*u*_1_ *>* 0 and *u*_2_ = 0).

### A. *Oct4 Overexpression: u*_2_ *>* 0*, u*_1_ = 0

#### 1) Disappearing ESC State Explains High Failure Rate

Fig. 5 summarizes the shapes of the modified nullclines for different values of Oct4 overexpression, *u*_2_. The chart in Fig. 5A shows three regimes for the number of SSS as a function of *u*_2_. There are 3 SSS for *u*_2_ ≈ 0 (Fig. 4A), 2 SSS for intermediate values of *u*_2_ (Fig. 5B), and 1 SSS for large values of *u*_2_ (Fig. 5C).

**Fig. 5:**
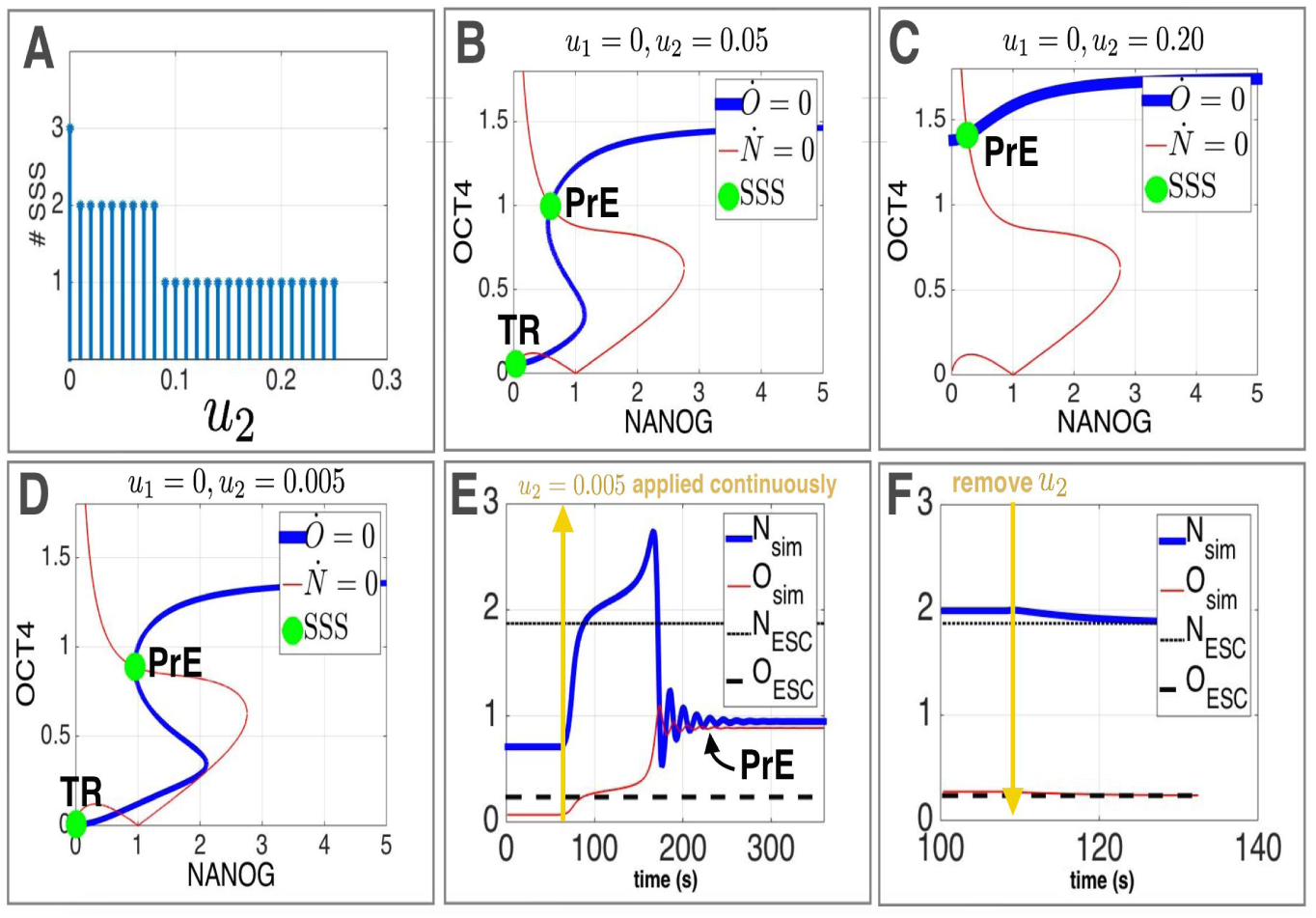
Oct4 overexpression summary (parameters adopted from Fig. 4): (A) Number of SSS as a function of Oct4 overexpression intensity, *u*_2_. (B) For *u*_2_ =0.05, the ESC state disappears. (C) For an even higher overexpression intensity, *u*_2_ =0.2, the TR and ESC states disappear, leaving only the PrE state. (D) For specific values of *u*_2_ ≈ 0.005, a near intersection occurs between the nullclines in the region where the ESC state originally was (Fig. 4A) (E) When *u*_2_ ≈ 0.005 (as in Fig. 5D) and a cell in the TR state is pushed out of its basin of attraction due to noise, it makes a transition to the PrE state. In the interim, it spends time at the near-intersection close to the original ESC state (Fig. 4A). (F) If Oct4 overexpression is stopped suddenly due to dilution or degradation in the beginning of the interim state in (E), the transitioning cell settles into the original ESC state, and is thus reprogrammed into an iPSC.

In Fig. 5B, a representative value of *u*_2_ from the intermediate range causes the 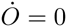 nullcline to change shape such that the intersection located at high Nanog, intermediate Oct4 – the region characterizing the ESC state – disappears. The TR and PrE states are still present and stable at this value of *u*_2_, so any cell starting at these states will never reach the ESC, which no longer exists.

In Fig. 5C, an even higher representative value of *u*_2_ causes the disappearance of the entire S-shaped region of the 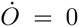 nullcline, which was creating the intersection corresponding to the TR state. As such, the system’s only remaining SSS is located in the PrE region (high Oct4, low/intermediate Nanog), so any cell starting in TR would be reprogrammed into the PrE state, hence missing the ESC state again.

These results support the idea that current reprogramming approaches have systematically low yields because constant overexpression shifts the nullclines in a manner to make the stable ESC state disappear, making it impossible to drive the system towards such a state.

#### 2) Extremely Rare Success

Nonetheless, [46] reports success – albeit at extremely low yields – in reprogramming TRs to ESCs with only Oct4. Our model 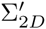 is able to replicate this unlikely success as demonstrated in Figs. 5E and 5F, which can be explained as follows. A narrow window of values around *u*_2_ ≈ 0.005 results in nullclines of the form seen in Fig. 5D, in which the 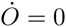 nullcline loses the hump in its shape enough to come very close to intersecting 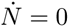 in the ESC region. This near intersection provides a window of opportunity for a cell to become an ESC.

In particular, if a cell is in the TR state and *u*_2_ ≈ 0.005, some intrinsic or extrinsic source of noise may push it out of the TR’s basin of attraction and closer to the neighboring unstable state (Fig. 5D). If this occurs, the cell could be repelled by the unstable state, and would ultimately transition to the PrE state as shown in Fig. 5E.

However, Fig. 5E also shows that en route to becoming a PrE, the cell is temporarily captured in a region very close to the original ESC state when *u*_2_*, u*_1_ =0 (Fig. 4A). Fig. 5F shows that if *u*_2_ approached zero during this time (due to dilution of the overexpressed Oct4), the cell settles to the original stable ESC state, and is thus reprogrammed from TR to ESC using only Oct4 overexpression.

The fact that successful reprogramming requires a specific value of Oct4 overexpression levels [37], [38], in addition to two other unlikely events occurring in series is one explanation for why overexpression-based reprogramming rarely succeeds in practice.

### B. *Nanog Overexpression: u*_1_ *>* 0*, u*_2_ = 0

Although we found no reports attempting to reprogram differentiated cells to ESCs via only Nanog overexpression, we perform an analysis on Nanog analogous to the one done above for Oct4 overexpression. We show that overexpression of Nanog also leads to disappearance of the ESC state.

#### 1) Disappearing ESC State Explains High Failure Rate

Fig. 6 summarizes the nearly equivalent results for Nanog overexpression as seen in Fig. 5 with Oct4 overexpression. Except for a certain range of values (*u*_1_ ≈ 0.12), the 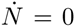 nullcline loses the hump in its shape, causing the disappearance of ESC states (Figs. 6B, 6C).

**Fig. 6:**
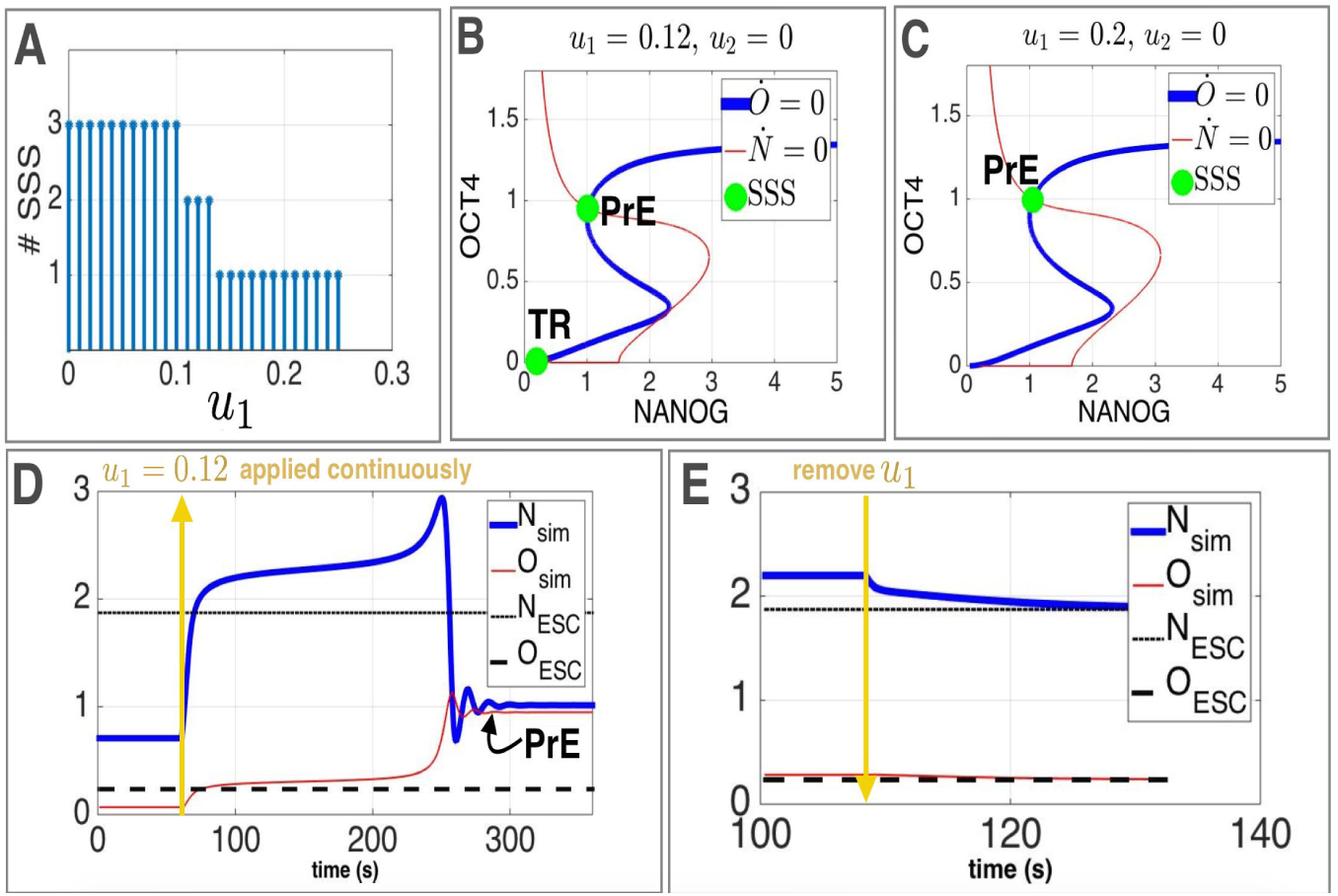
Nanog overexpression summary (parameters adopted from Fig. 4): (A) Number of SSS as a function of Nanog overexpression intensity, *u*_1_. (B) For *u*_1_ = 0.12, the ESC state disappears. A near intersection between the nullclines still occurs next to where the ESC state once was (Fig. 4A) (C) For an even higher overexpression intensity, *u*_1_ = 0.2, the TR and ESC states disappear, leaving only the PrE state. (D) When *u*_1_ ≈ 0.12 (Fig. 6B) and a cell in the TR state is pushed out of its basin of attraction due to noise, it makes a transition to the PrE state. In the interim, it spends time at the near-intersection close to the original ESC state (Fig. 4A). (E) If overexpression is stopped suddenly due to dilution or degradation in the beginning of the interim state in (D), the transitioning cell settles into the original ESC state, and is thus reprogrammed into an iPSC.

#### 2) Extremely Rare Success

As seen in Fig. 6D, for values of *u*_1_ ≈ 0.12, a TR cell that is pushed out of it’s basin of attraction is then repelled by the nearby unstable state, and can be pushed towards the PrE state. Fig. 6D shows that en route to the PrE state, the cell is temporary captured in a region near the original ESC state when *u*_2_*, u*_1_ = 0. If Nanog overexpression is removed during this temporary suspension (*u*_1_ = 0), the cell settles to the stable ESC state and is thus reprogrammed into an iPSC (Fig. 6E).

Although many reprogramming experiments involve the use of multiple transcription factors, the systematic failures demonstrated here with single-factor overexpression can be seen to carry through if both Oct4 and Nanog were overexpressed at the same time (*u*_1_ *>* 0*, u*_2_ *>* 0). In fact, in this case the nullclines would intersect for *u*_1_ = *u*_1_ = 0.2 at points given by the intersection of 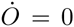 in Fig. 5C and 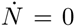 in Fig. 6C. It can be seen by inspection that these two nullclines intersect only once, in the PrE region.

## IV. Conclusion

In this work, we have modeled the difficulty of iPS cell reprogramming using standard tools from dynamical systems theory. Our study was based on understanding transitions among basins of attraction in a multi-stable system under additive positive inputs. It is very well known that for some bistable systems such as the toggle switch [47] or its variants [48], one can always force transitions from any initial state to any desired basin of attraction by suitable applications of positive additive input. In contrast to this, the structure of the network studied in this paper does not necessarily allow such forced transitions.

In future work, we will study the general principle that makes the distinction between networks in which such transitions can be forced with positive additive inputs, and those in which this is not possible. We will also investigate theoretical overexpression models that use closed-loop feedback to overcome this structural problem.

## Notes

* This work was supported by the Paul E. Gray Fund and AFOSR-BRI grant #FA9550-14–1-0060.

